# Topography and Land Use Modulate Hydrological and Nutrient Export Dynamics in two Andean Streams

**DOI:** 10.1101/742411

**Authors:** Alejandro Sosnovsky, Mailén Elizabeth Lallement, Magalí Rechencq, Eduardo Enrique Zattara, María Valeria Fernández, Sergio Leiva, María José Suárez, Romina Daga, Rodolfo Juan Carlos Cantet

**Affiliations:** Grupo de Evaluación y Manejo de Recursos Ícticos, INIBIOMA, CONICET-Universidad Nacional del Comahue, Quintral 1250, R8400FRF San Carlos de Bariloche, Argentina; Grupo de Nuevos Materiales y Dispositivos, Comisión Nacional de Energía Atómica Av. Ezequiel Bustillo Km 9.5 (8400) San Carlos de Bariloche, Argentina; Departamento de Producción Animal, Facultad de Agronomía, Universidad de Buenos Aires, e INPA, CONICET. Av. San Martín 4453, C1417DSQ Ciudad Autónoma de Buenos Aires, Argentina; Laboratorio de Análisis por Activación Neutrónica, Centro Atómico Bariloche, Comisión Nacional de Energía Atómica, Av. Bustillo 9500, 8400 Bariloche, Argentina Centro Científico Tecnológico CONICET. Patagonia Norte.

**Keywords:** Patagonia, Flashiness, Autocorrelation, Cross-correlation, Discharge, Nitrogen, Phosphorus, Eutrophication

## Abstract

Fluvial dynamics are driven by multiple environmental factors and scales. Studies coupling hydrological and nutrient dynamics of Andean streams are almost nonexistent. We characterized two adjacent streams with contrasting drainage basins: Casa de Piedra, originating in a small mountain lake and running through a pristine landscape, and Gutiérrez, originating in a large piedmont lake and running through an anthropized landscape. Despite both drainage basins sharing geology and climate, we found that the streams presented contrasting hydrological and nutrient dynamics. Casa de Piedra had higher discharge flashiness with shorter response delays to precipitation. Interestingly, Gutiérrez’s hydrology was buffered by the upstream lake, but its nutrient exports were not. Differences observed in quality and timing of coarse particulate organic matter export and basal export levels of phosphorus and nitrogen could be explained by human activities affecting Gutiérrez but not Casa de Piedra. Moreover, nitrogen:phosphorus ratio indicates a possible future shift to phosphorus as the limiting nutrient as Andean basins become more densely populated. In summary, our annual basis study shows that even under a common geology and climate, stream dynamics of adjacent basins can be starkly different due to differences in topography and land use.

## INTRODUCTION

River ecosystems have traditionally been studied as isolated entities; however, it has recently become clear that they are deeply intertwined with their surroundings. Climate, topography, geology and land cover are environmental factors that show a hierarchical influence on spatial patterns of these ecosystems (Snelder and Biggs 2002). In other words, a river is the product of its landscape (Wiens 2002) and thus riverine hydrological, chemical and biological processes should be studied in relation to the river’s overall drainage basin (Hynes 1975).

Regional climate and drainage basin topography are factors acting at a macro and meso spatial scales upon the river ecosystem (Snelder and Biggs 2002). Climate directly drives the hydrological cycle, determining the flow regime of a stream, a key ecosystem variable (Poff et al. 1997). Rain, snowmelt and groundwater feed stream discharge, the relative importance of these factors varying over time and space (Brown et al. 2003). After a rain episode, surface and sub-surface run-off are favored at mountainous landscapes, while water infiltration and evapotranspiration are favored at highly vegetated lowlands with permeable soils. Snowfall at high altitudes contributes to formation of a snowpack, which will store water until the melt period, when this water flows down into a stream (Bailey 1995). Lakes are topographic features of a drainage basin that can buffer run-off response and sediment transport (Snelder and Biggs 2002). Thus, headwater lakes influence directly the hydrograph and nutrient dynamic of downstream fluvial ecosystems (Gordon et al. 2004). These hydrographs tend to be stable over time, furthermore, considering lakes as sediment and nutrient traps, low values of nutrient concentration and export could be expected in these streams (Little et al. 2008; Parker et al. 2009).

At a lower spatial scale, drainage basin traits also affect fluvial dynamics, since nutrients and contaminants generated at the basin eventually reach the river ecosystem (Allan 2004; Dodds and Smith 2016). Non-point source pollution, associated with changing land use patterns and practices, has resulted in increased impacts on water bodies (Jordan et al. 1997; Carpenter et al. 1998). For example, growth of urban areas favors nutrient export (Howarth et al. 1996; Meyer et al. 2005; Walsh et al. 2005). Moreover, direct dumping of sewage water to a stream greatly increases nutrient load (Pieterse et al. 2003) and can also change nutrient stoichiometry (Merseburger et al. 2005). Thus, increases of nutrient load leading to eutrophication of running waters and carbon dynamics largely depend on the drainage basin state.

In Northern Patagonia, characteristics of Andean ecosystems are largely determined by the Andes mountain range. This region is characterized by a temperate-cold climate with warm and dry summers (Peel et al. 2007). The Andes represent an important barrier for humid air masses brought from the Pacific Ocean by westerly winds. Most of the humidity in these maritime air masses precipitates on the west side of the Andes as they are blown to higher, colder altitudes across the mountain range. Upon crossing the Andes, the air descends along the east slopes and becomes hotter and drier through adiabatic warming. East of the Andes, the amount of rainfall follows a steep west-east gradient: annual precipitation drops from 3500 mm to 700 mm in less than 60 km. Moreover, the precipitation regime shows marked seasonality, with most rain falling during fall and winter (Paruelo et al. 1998). As in other South American watersheds, N deposition is remarkably low (Holland et al. 1999), and forest (Diehl et al. 2003) and lake ecosystems (Diaz et al. 2007) are limited mainly by N. Moreover, a large fraction of drainage basins along the East side of the Andes are in a relatively pristine state because human population density is low. Diverse authors have studied anthropic (Miserendino et al. 2016) and natural (Lallement et al. 2016; Carrillo et al. 2018; Williams Subiza and Brand 2018) impacts on Andean drainage basins and their fluvial ecosystems. Recent research has revealed that dynamics of organic matter are intimately associated with the stream-basin interaction (García RD et al. 2015; Díaz Villanueva et al. 2016; García et al. 2018). Andean streams originate mainly from snowmelt, glacial melt and piedmont lakes and are predominantly oligotrophic (Pedrozo et al. 1993). Sosnovsky et al (2020) described their discharge and physico-chemical dynamics during the three contrasting hydrological periods characteristics of mountain regions, storm, snowmelt and basal periods. Oyarzún et al (2004) and Temporetti (2006) estimated the nutrient load of pristine and burned watersheds, respectively. Nevertheless, studies coupling hydrological and nutrient dynamics in the region are lacking, despite the hydrological, ecological and human importance of these variables and their possible interactions.

The main hypothesis of our study is that climate is the main driver of stream ecosystems dynamics, which are in turn modulated by topography and land use. Therefore, we expect precipitation, air temperature and stream discharge to be intimately related. Under this hypothesis, adjacent drainage basins with contrasting topography and land use should present differences in their hydrological, physical and chemical dynamics. To test this expectation, we selected two streams located in adjacent yet contrasting drainage basins: Casa de Piedra (CP) and Gutiérrez (G). CP stream originates in a small high-mountain lake and its drainage basin is pristine throughout its length. In contrast, G stream originates from a large piedmont lake which buffers it hydrological dynamic (Sosnovsky et al. 2020), and its drainage basin suffers from significant anthropization. While the large headwater lake is expected to buffer chemical dynamics in G, higher anthropization of G’s basin could impact the stream ecosystem, potentially causing increase nutrient concentration and export. Our first goal was to characterize discharge dynamic within a year in these fluvial ecosystems. To this end, we compare discharge magnitude and variability between CP and G, and study how it relates to climatic variables (basin precipitation and air temperature). Our second goal is to determine the trophic state of both streams, measuring basal concentration, export and stoichiometry of nitrogen (TN: total N) and phosphorous (TP: total P), and estimating basal export of coarse particulate organic matter (CPOM, organic matter > 1mm). Our third goal is to identify the seasonal patterns of key physical and chemical variables.

## MATERIALS Y METHODS

### Study area

Regional landscapes of Andean North Patagonia are the result mainly of glaciation/deglaciation, fluvial and mass movement processes over the previous structural-tectonic landscape. In the region, most fluvial courses have clear structural control. Highest mountains show mainly the result of glacial erosion processes while both erosive and depositional geoforms are present in the hillsides corresponding to ancient lateral glacial valleys. The alluvial plains of streams and tributaries in the area run mainly on moraine systems and stream mouths form alluvial fans over glacifluvials plains (Pereyra et al. 2005).

The lithology of the study site is mainly composed of rocks of Paleozoic metamorphic and granitic basement, Cretacic and Miocene granite porphyry, and limited outcrops of Jurassic volcanic and sedimentary rocks covering previous granitoids. Oligocene volcanic and marine sedimentary rocks are also present (Giacosa et al. 2001). In general, the rocky outcrops constitute the hydrogeologic basement with almost absent primary permeability and secondary permeability acquired by folding and fracturing of the rocks. Headwaters are dominated by rocky outcrops and poor soil development is associated mainly to presence of vegetation and lower slopes on the sides of glacial valleys. On the other hand, the lower parts of the landscape are represented by Pliocene-Pleistocene glacial material (moraines, glaciofluvial-fluvial and glaciolacustrine deposits) with poorly to moderately developed soils (Andisols reaching more than 80 cm in some places) under well drained conditions, well provided of superficial organic matter and phosphorus retention >85% (Satti et al. 2003). The most recent sedimentary deposits correspond to the present alluvial materials (gravel, sand, silt) in the flood plains surrounding the Gutiérrez lake and streams, with poorly developed soils due to active morphodynamics and coarse materials (Giacosa et al. 2001; Pereyra et al. 2005). Associated to these deposits are unconfined aquifers of variable permeability and discontinuous character due to the heterogeneity on the grain size of the deposits (Pereyra et al. 2005).

Research was conducted at the drainage basin of the streams Casa de Piedra (CP; 41° 07’ 30.11” S 71° 27’ 13.16”W) and Gutiérrez (G; 41° 09’ 36.18” S 71° 24’ 37.19”W), both located in Nahuel Huapi National Park, Río Negro, Argentina (Figure 1). These drainage basins are very different from each other, despite of their proximity (Table 1). CP’s basin covers 65.5 km^2^ and the stream originates in lake Jakob, a small (0.15 km^2^), ultra-oligotrophic (1-3 μg l^−1^ TP), high altitude water body situated at 1550 m above sea level (García P et al. 2015). CP’s only perennial affluent is the Rucaco stream, which originates at a high-altitude wetland. CP flows 19.7 km through a steep V-shaped valley (average slope: 33.9 m km^−1^) for most of its length. CP’s drainage basin is mostly pristine, with human activities restricted to hiking and infrequent camping; human population (less than 1000 inhabitants, 2010 National Census) is restricted to lowland areas. Forest below the tree line includes deciduous *Nothofagus pumilio* and evergreens *N. dombeyi* and *Austrocedrus chilensis*. In contrast, G’s drainage basin is larger (161.6 km^2^), and the stream originates in lake Gutiérrez, a deep (111 m maximum depth) and large (17 km^2^) water body that occupies ∼10 % of the basin’s area. The seasonal concentration of TP is 3.4 μg l^−1^ on average (Diaz et al. 2007). G’s only perennial affluent is the Cascada stream, whose catchment covers 13.0 km^2^ and includes a major winter sports resort. G flows 9 km through a wide, gently sloped valley (5.9 m km^−1^). Its riverbanks are extensively colonized by the exotic crack willow, *Salix fragilis*, and pass through a series of populated areas (approximately 4500 inhabitants, 2010 National Census) belonging to the city of San Carlos de Bariloche (approximately 108000 inhabitants, 2010 National Census). In addition, one trout farm with a total production of ∼10 tons year^−1^ use G’s waters. In summary, CP’s basin is largely free of impact from human activities, while G’s basin is not.

**Figure 1.**
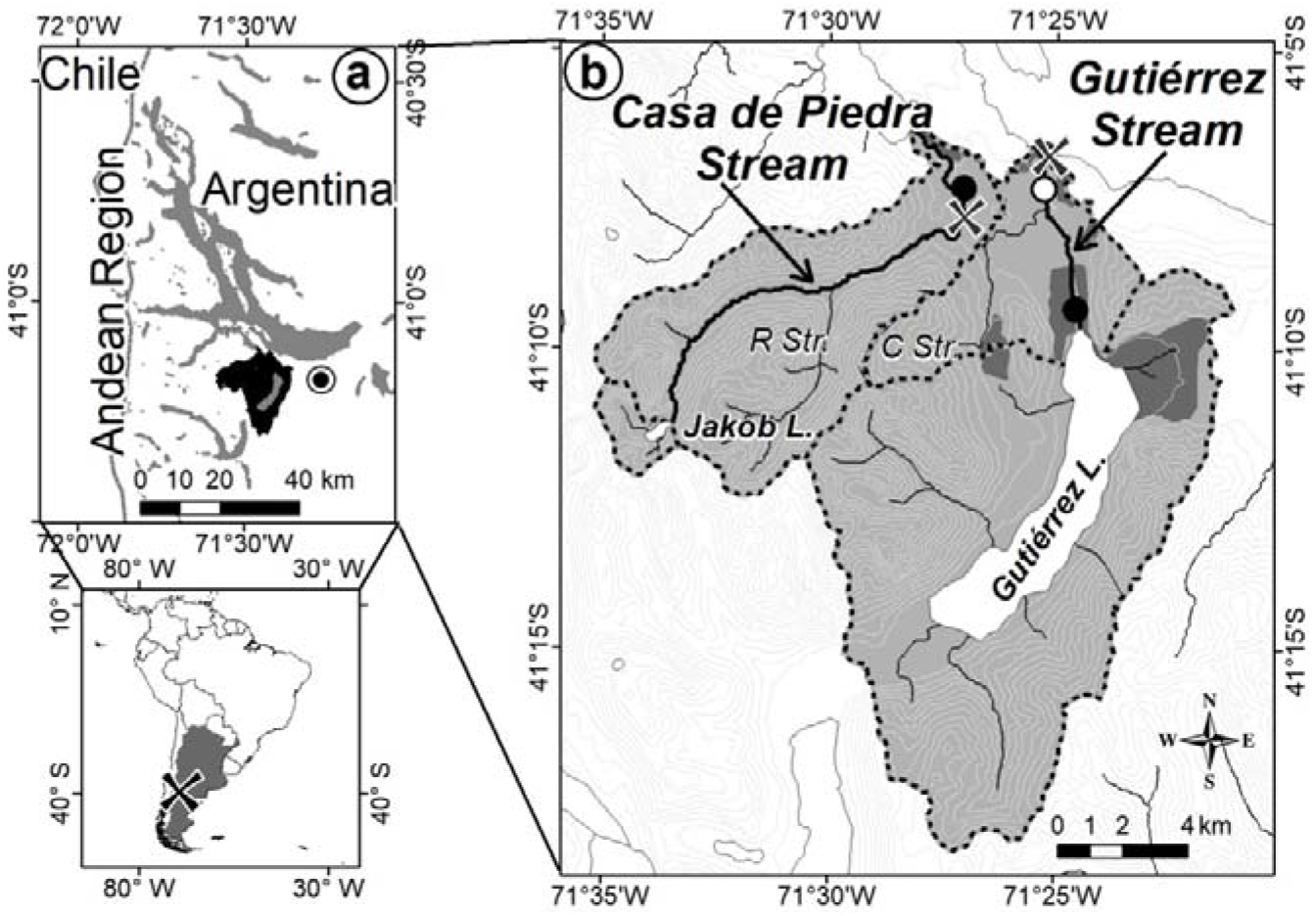
a) Nahuel Huapi National Park; the study site is shaded in black. The city marker indicates San Carlos de Bariloche downtown. b) Study area, Casa de Piedra and Gutiérrez drainage basins, Jakob lake and Gutiérrez lake sub-catchments are also delimitated. Curves represent 100 m slope on the terrain. Populated zones are shown in dark grey, crosses indicate sampling sites, black circles indicate the limnigraphic stations and the white circle indicate the rain meter location. R Str: Rucaco stream; C Str: Cascada stream (with permission from Sosnovsky et al. 2020).

**Table 1.** Drainage basins characteristics. Values are expressed based on the entire area of the drainage basins, values between parentheses exclude the sub-catchment areas draining into Jakob lake and Gutiérrez lake.

|  | **CP:** Casa de Piedra | **G:** Gutiérrez |
| --- | --- | --- |
| Drainage area (km<sup>2</sup>) | 65.5 (60.3) | 161.6 (30.1) |
| Lake area (km<sup>2</sup>) | 0.15 | 17.21 |
| Native forest (%) | 60.2 (61.7) | 61.1 (56.3) |
| Bare ground (%) | 29.7 (27.7) | 17.2 (9.5) |
| Exotic forest (%) | 9.9 (10.5) | 10.8 (34.2) |
| Population (N) | < 1000 (< 1000) | 4528 (3580) |
| Urban area (km<sup>2</sup>) | 0.65 (0.65) | 11.35 (5.08) |

### Data collection

The present study was carried out over a period of one year, from March 2013 to March 2014. Daily precipitation was recorded with a rain meter (Figure 1) and did not differentiate between rain and snow. Discharge data were taken from average daily records from limnigraphic stations. CP’s limnigraphic station is located close to its mouth, while G’s station is located at its source (Figure 1). Values of discharge at our G sampling site were estimated as discharge as measured at the station plus discharge for the Cascada stream, determined using the Drainage-Area Ratio method (Emerson et al. 2005); based on the specific daily values of discharge for the adjacent CP stream. Specific discharge (Q^sp^) was defined as discharge per unit area. Daily average air temperature records were obtained from the National Weather Service.

Baseline levels of nutrients and relevant physical and chemical variables were obtained from monthly samples at both streams (Figure 2). Water samples were taken to analyze total phosphorus (TP) and total nitrogen (TN) concentrations. Water turbidity was measured with a Velp turbidimeter. Temperature, electrical conductivity (EC) at 25°C and pH were measured with an Oakton probe. Coarse particulate organic matter (CPOM) was sampled with 2 drift nets (1-mm, 20 × 20-cm opening at the mouth) that were staked in the streambed and left to collect material for intervals of 10 to 200 minutes, depending on the amount of CPOM present in the stream. During October, high and fast stream discharge prevented us from placing Surber sampler nets at G, so we performed linear interpolation for this value. Data were taken exclusively on days without precipitation.

**Figure 2.**
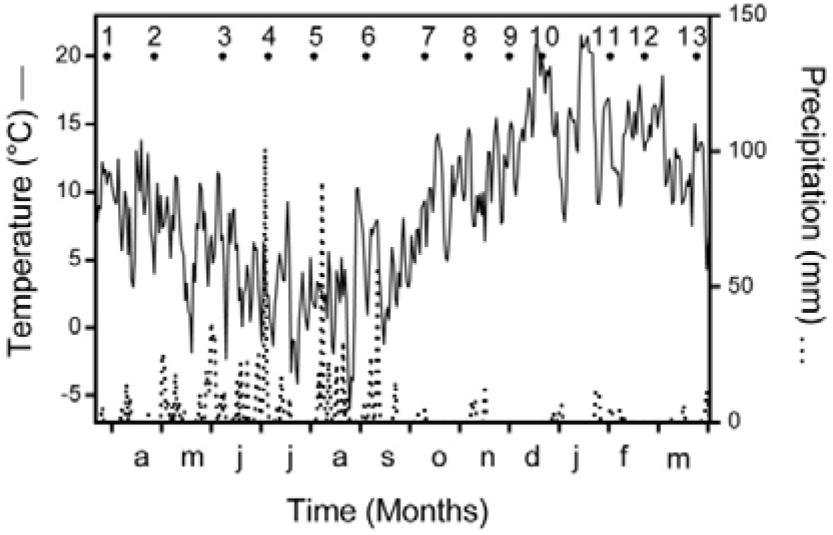
Temperature (solid line) and precipitation (dotted line) during the study period. Black dots and numbers indicate sampling dates.

### Data processing and analysis

#### Discharge

Differences in stream discharge were evaluated using a heterogeneous variance model. For this purpose, data on the average daily discharge and average daily specific discharge were transformed to normality using the Box-Cox procedure as suggested by Peltier et al. (1998). To understand the dynamics of discharge, we used an autocorrelation analysis (Mangin 1984). This analysis examines how a value depends on preceding values over a period of time. The correlations are computed for a given lag time k to obtain an autocorrelation coefficient r(k). For example, the correlation between today’s and yesterday’s stream discharges would be a lag-1 autocorrelation coefficient, r(1). The autocorrelation function is represented with a correlogram, where slope is determined by the response of the system to an event. If the event has only a short-term influence on the response of the discharge, the slope of the correlogram will decrease steeply. In contrast, if the system is influenced by an event for a long time, the slope of the correlogram will decrease slowly. Generally the length of the influence of an event is given by the “memory effect” which is according to Mangin (1984) the lag number k when r(k) reaches a value of 0.2. The formula for autocorrelation is (Larocque et al. 1998):

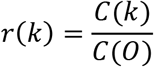

with

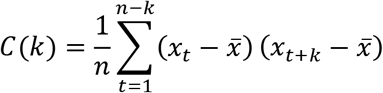

Where k is the time lag and varies from 0 to n. According to Mangin (1984), n has to be taken as 1/3 of the whole dataset to avoid stability problems.

In addition, we studied the relationship between discharge, precipitation and temperature by cross-correlation analysis. This analysis is widely used to analyze the linear relationship between input and output signals in hydrology (Larocque et al. 1998). In this case, input signals were precipitation and temperature, and output signal was river discharge of CP and G streams. Cross-correlations are represented by a cross-correlogram. The maximum amplitude and the lag value of the cross-correlogram provide information on the delay, which indicates the time of the pulse transfer to the stream. According to Larocque et al. (1998) the formula for cross-correlation is:

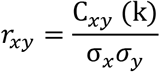

with

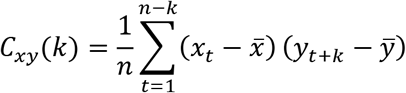

Where σx and σy are the standard deviation of the two times series.

#### Physical and chemical analysis

We estimated concentration and export of nutrients (TN and TP) and CPOM. TN was analyzed through digestion and subsequent reduction in cadmium column (Grasshoff et al. 1983). TP was analyzed using acid digestion followed by evaluation of solubilized phosphorus as soluble reactive phosphorus (A.P.H.A. 2005). We estimated Ash Free Dry Weight of the CPOM using the Pozo et al. (2009) methodology. The export of TN, TP and CPOM were calculated for each sample date by the following formula:

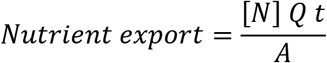

Where [N] is instantaneous nutrient concentration (mg m^−3^), Q is instantaneous discharge (m^3^ s^−1^), *t* is time (yr) and A is drainage basin area (km^2^). The annual nutrient export was an average of each data set (n=13).

We performed a paired T-test to compare nutrient concentration and export between streams; to this end the data were transformed to comply with normality and homoscedasticity assumptions. Last, we performed a Principal Component Analysis (PCA) that included 12 variables: turbidity, temperature, EC, pH, discharge, TP, TN, TN:TP, CPOM, TP export, TN export and CPOM export. PCs were calculated based on standardized correlation matrix, using components with eigenvalues larger than 1. Number Cruncher (Hintze 1998) was employed for statistical analysis and GraphPad Prism version 6.01 was employed for graphics editing.

## RESULTS

### Climate and hydrology of the region

Air temperature was low between May and August and increased monotonically after September, reaching its highest values from December to February. Annual precipitation was 1164 mm and showed strong seasonality, with 80% falling between mid-May and mid-September. The first two (#1-2) and last three (#11-13) samples thus took place during a drier period; samples #3-6 were taken during the precipitation period; the remaining samples (#7-10) fell within the period of monotonic air temperature warming (Figure 2). Thus, samples were evenly distributed across three climatically contrasting periods.

Cross-correlation of discharge with precipitation and temperature showed opposing patterns. Discharge was positively correlated with precipitation, but negatively correlated with air temperature (Figure 3c, d, Appendix S1). The discharge-precipitation correlogram shows three periods that coincide with peaks of the correlation function (Figure 3c). Both streams show differences at each of these peaks: CP had higher correlation values (r max) and lower lag times than G. Indeed, at CP the correlation reached its maximum the same day precipitation took place (lag = 0) during the first period, while r was not even significant for G at the time (Appendix S1). During the second period, correlation at CP reached an r max=0.36 with a lag time of 34 days, while at G, correlation was lower (r max=0.19) and the lag higher (lag = 40). The third, longest period presented a series of peaks and valleys, and lapsed from day 57 to day 113 after the precipitation event at CP and from day 62 to 145 at G. In contrast, the discharge-temperature correlogram did not show similar periods; cross-correlation was significant from day 0 to around day 90 for both streams (Figure 3d, Appendix S1).

**Figure 3.**
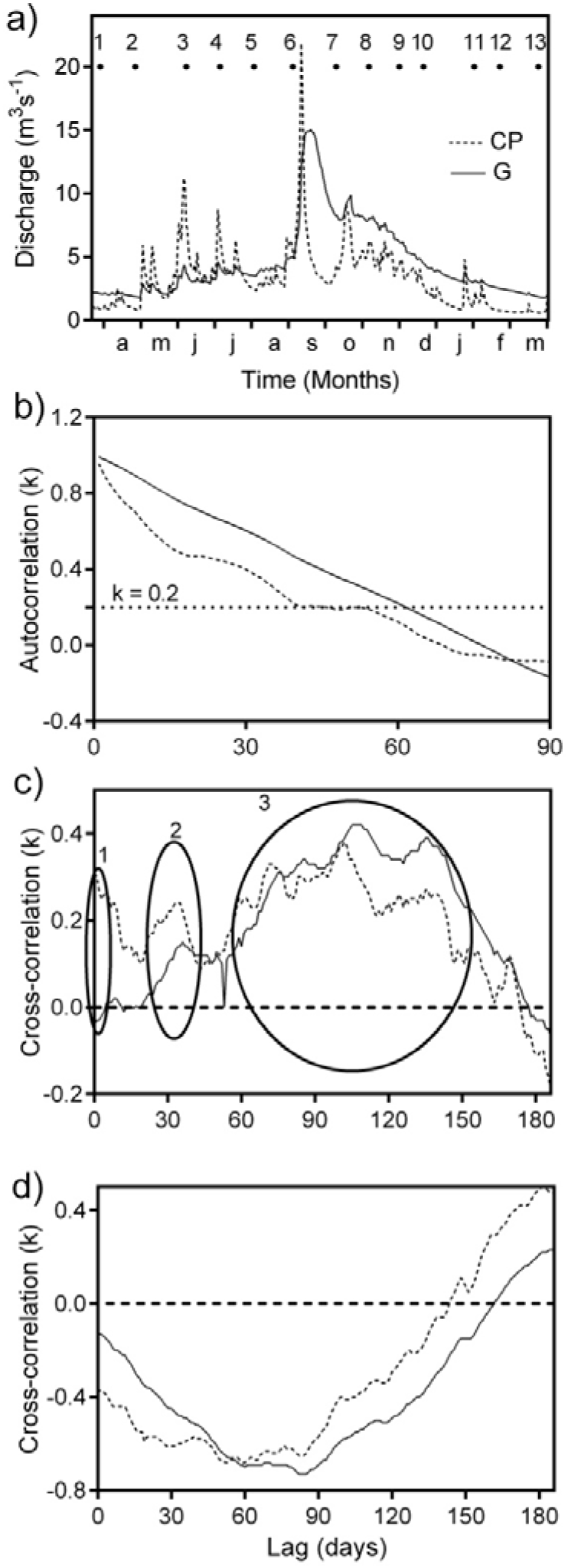
a) Hydrographs of Casa de Piedra (CP) and Gutiérrez (G) streams, black dots and numbers indicate sampling dates. b) Autocorrelation functions and memory effects (r (k) = 0.2) of the discharge. c) Cross-correlations between discharge and precipitation, three different periods are shown, and d) cross-correlations between discharge and temperature.

The flow regime was contrasting between the streams. CP had a lower annual average discharge value than G (Discharge CP= 3.364 m^3^ s^−1^; Discharge G= 4.635 m^3^ s^−1^; t =7.4437; P < 0.0001; d.f. = 728), but its Q^sp^ was higher (Q^sp^ CP= 0.051 m^3^ s^−1^; Q^sp^ G= 0.029 m^3^ s^−1^; t=−24.7917; P < 0.0001; d.f. = 728). CP presented the most extreme discharge values (Figure 3a) and a steep autocorrelation function with a memory effect at k = 38, while G revealed a more gently sloped function with a memory effect at k = 61 (Figure 3b). Thus, CP showed lower memory effect than G, indicating a higher flashiness of stream discharge.

### Physical and chemical characteristics of the streams studied

Although streams differed in the specific values, both had very low EC, turbidity, nutrient concentration and nutrient export (Table 2). G showed the highest TP and TN concentrations (Figure 4a, c) and exports (Figure 4b, d): CP exported 8.6 kg TP km^−2^ yr^−1^ and 57.5 kg TN km^−2^ yr^−1^, while G exported 11.6 kg TP km^−2^ yr^−1^ and 78.4 kg TN km^−2^ yr^−1^ (approximately 35 % more P and N). No significant differences in TN:TP average ratios were found (∼15; Figure 4e), although the median value of TN:TP was 11 for CP and 16 for G. As to the amount of CPOM exported, no significant difference between streams was found. There were differences, however, in the quality of exported material; native *N. pumilio* leaves dominated in CP, while introduced *S. fragilis* leaves dominated in G. The peaks of CPOM export also differed among streams: highest export at CP was found during June and July (samples #3-4), while at G this peak occurred earlier, in April (sample #2). In summary, the drainage basin showing more anthropic land uses also showed higher nutrient export, with most CPOM export represented by introduced tree species.

**Table 2.** Average values, maximum and minimum in brackets, of the physical and chemical variables in the streams (n=13, except for the Coarse particulate organic matter where n=12).

|  | Casa de Piedra | Gutiérrez |
| --- | --- | --- |
| Temperature (°C) | 7.8 (3.7-12.5) | 10.6 (6.0-16.8) |
| Electrical Conductivity<br>(µs cm<sup>-1</sup>) | 43 (27-61) | 74 (71-76) |
| Turbidity (NTU) | 0.2 (0.0-0.8) | 0.7 (0.0-2.0) |
| pH | 7.8 (7.4-8.0) | 7.9 (7.3-8.1) |
| Total Phosphorus (mg m<sup>-3</sup>) | 5 (2-9) | 14 (10-18) |
| Total Nitrogen (mg m<sup>-3</sup>) | 29 (18-44) | 94 (75-134) |
| TN:TP (molar) | 15 (6-31) | 15 (11-21) |
| Coarse Particulate Organic<br>Matter (AFDW m<sup>-3</sup>)*10 | 0.46 (0.04-1.86) | 0.91 (0.02-8.76) |
| Total Phosphorus export<br>(kg km<sup>-2</sup> yr<sup>-1</sup>) | 8.6 (1.7-25.4) | 11.6 (3.8-22.5) |
| Total Nitrogen export<br>(kg km<sup>-2</sup> yr<sup>-1</sup>) | 57.5 (7.9-129.6) | 78.4 (27.3-184.8) |
| Coarse Particulate Organic<br>Matter export (kg km<sup>-2</sup> yr<sup>-1</sup>) | 126.4 (1.3-750.4) | 43.1 (1.5-317.9) |

**Figure 4.**
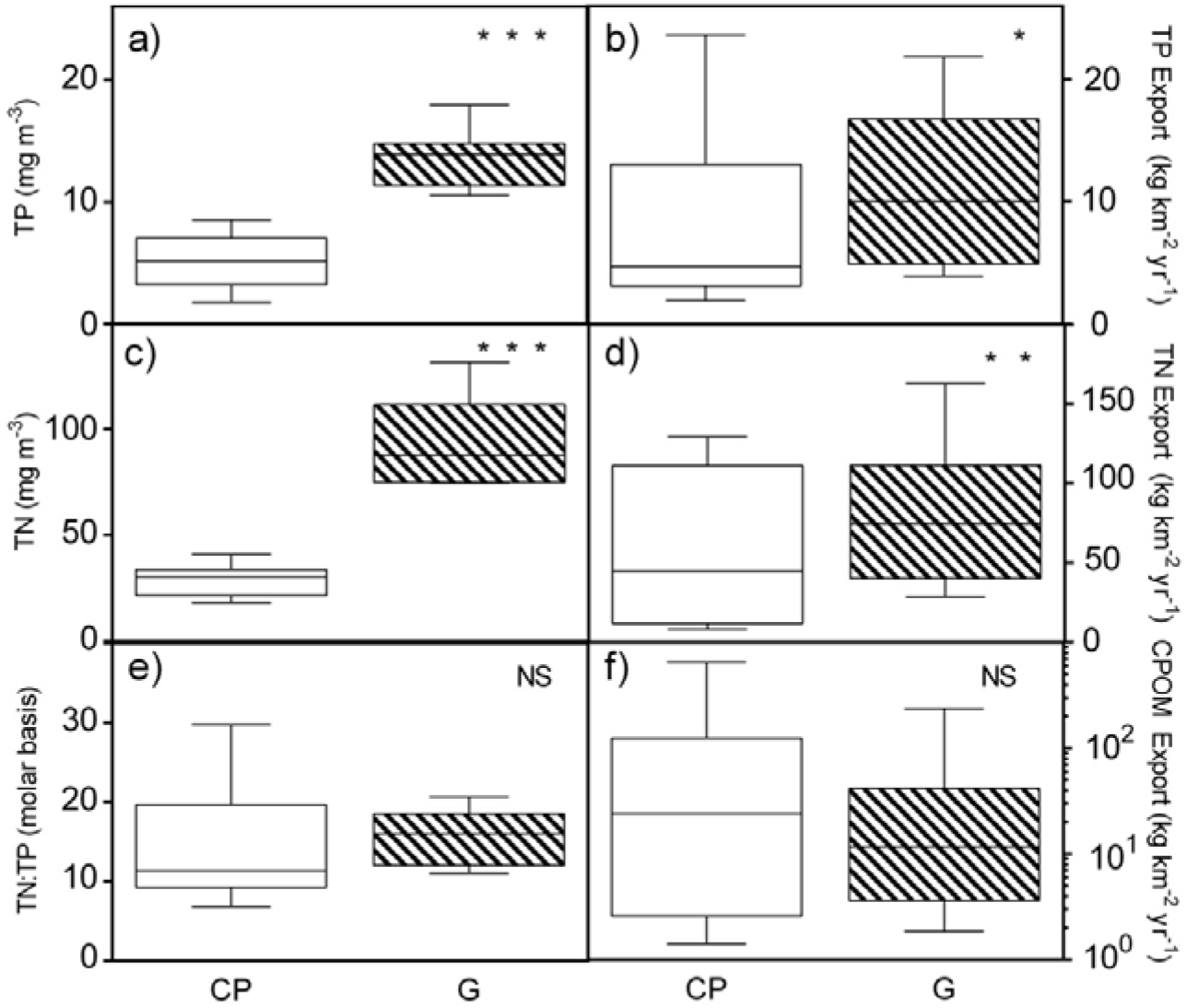
Box plot showing annual differences in nutrient concentration, ratio and export in Casa de Piedra (CP) and Gutiérrez (G) streams (Paired T Test, * P ≤ 0.05; * * P ≤ 0.01; * ** P ≤ 0.001; ns: not significant). f) The Y axis is in logarithmic scale. Total Phosphorus (TP), Total Nitrogen (TN) and Coarse Particulate Organic Matter (CPOM).

Our PCA determined three Principal Component, FI, FII and FIII that respectively explain 33%, 24% and 13% of the variance, accounting for 70% of the total variance (Table 3). FI was mainly related to discharge and nutrient export (TP_exp_ and TN_exp_) (Figure 5a). Positive FI values corresponded to samples with high discharge, high nutrient export and high turbidity. The FII was mainly related to the EC of the water, and to a lesser extent with water temperature and nutrient concentration (Figure 5a). The FIII was mainly related to the TN:TP ratio, and to a lesser extent with CPOM variables.

**Table 3.** Principal Component Analysis on physical and chemical variables. Factors are a linear combination of the different variables. Percent of variance explained and factor loadings (> 0.30) are shown. Abbreviations are as in Figure 5.

| Variable | First Factor<br>(33%) | Second Factor<br>(24%) | Third Factor<br>(13%) |
| --- | --- | --- | --- |
| Temperature |  | 0.31 |  |
| EC |  | 0.47 |  |
| Turbidity | 0.35 |  |  |
| Q | 0.38 |  |  |
| TP | 0.35 | 0.41 |  |
| TN | 0.36 | 0.32 |  |
| TN:TP |  |  | 0.55 |
| CPOM |  |  | -0.47 |
| TP<sub>exp</sub> | 0.42 |  |  |
| TN<sub>exp</sub> | 0.42 |  |  |
| CPOM<sub>exp</sub> |  | -0.35 | -0.55 |

**Figure 5.**
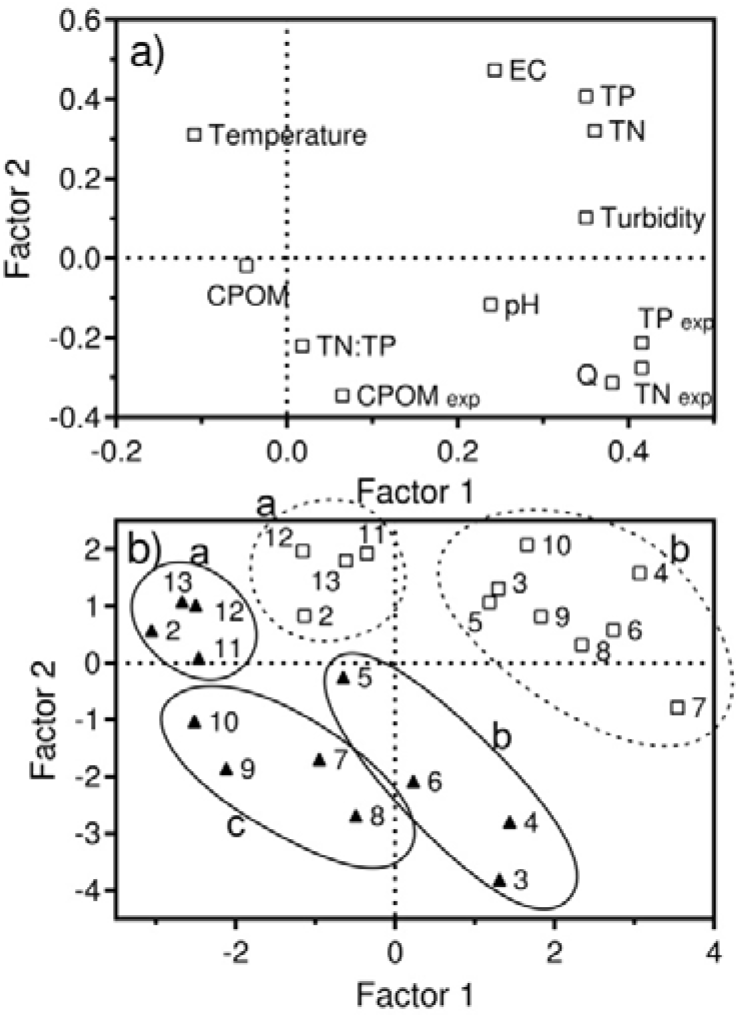
Principal Component Analysis (PCA) of physical and chemical variables in the study streams. a) The weight of variables in each of the axes is shown. b) First factorial plane of the PCA, Casa de Piedra sampling dates are represented by triangles and Gutiérrez ones are represented by rectangules, the numbers correspond to the sampling dates and they are grouped by climatic characteristics of the region (a, b and c). Electrical conductivity (EC), Discharge (Q), Total Phosphorus (TP), Total Nitrogen (TN), Coarse Particulate Organic Matter (CPOM), Export (exp).

Both streams are clearly discriminated by PCA ordering, highlighting seasonal differences in each stream (Figure 5b). Besides separating samples from each stream, plotting FI versus FII shows that each stream is temporally heterogeneous, allowing clustering in 3 groups for CP (CPA, CPB and CPC) and 2 for G (GA and GB) (Figure 5b). CPA had low discharge and export rate values and high temperatures, taking place at the warmest period (Figure 2). CPB had higher nutrient concentration and export values, matching the period with more abundant precipitation (Figure 2). CPC was mainly characterized by low EC, nutrient concentration and turbidity values, and coincides with the period of diminishing rainfall and monotonic air temperature warming (Figure 2). GA had similar characteristics to CPA and spans the same period. The remaining samples fall within GB. Thus, the values of physical and chemical variables sampled monthly through the year show that both streams have marked seasonal differences that follow precipitation and air temperature patterns.

## DISCUSSION AND CONCLUSIONS

We surveyed discharge and nutrient dynamics in two adjacent North Patagonian streams with contrasting drainage basins. Precipitation had a direct influence on stream discharge at both streams, but response time and amplitude varied between them. CP stream was flashy, showed a tighter relation between precipitation at the basin and discharge response and stronger seasonality of physical and chemical parameter values. G stream, in contrast, showed a more buffered discharge response to precipitation and higher nutrient exports. In general, similarities seen between streams can be related to regional climate and geology, while differences are better explained by contrasting topography and land use of the respective drainage basins.

Precipitation and temperature have opposite influence in river discharge in the Andean region (Masiokas et al. 2008). Precipitation increased stream discharge of our study sites. Run-off, groundwater and snowmelt, the different pathways taken by precipitated water to reach the streams and their relative importance are evidenced by the three periods found in the discharge-precipitation correlogram. Surface run-off was relevant at CP, its basin characterized by steep slopes. In this stream, discharge increased drastically during the same day precipitation occurred, and then during the following days. On the other hand, surface run-off was not relevant at G stream, likely due to buffering by the large upstream lake. However, at a shorter temporal scale, Sosnovsky et al. (2020) reported a positive relation between G stream discharge and precipitation with a 4-days lag during the storm season. Water not reaching the stream immediately as run-off infiltrates into the soil and moves to the stream as sub-surface run-off or groundwater; these pathways usually result in delayed stream discharge responses (Kalff 2002). Sub-surface run-off and groundwater pathways were more relevant at CP, which showed a higher r(max). However, delay time was similar for both streams; this is likely due to shared basin characteristics driving groundwater dynamics: lithology, geomorphology and soil development (Giacosa et al. 2001). In colder regions, snow precipitation is stored during the winter months and released as snowmelt in spring and summer (Bailey 1995). This results in a delay time between precipitation and discharge response of about 2-4 months, as we found at both study sites during spring. The observed discharge dynamics and its relationship with precipitation (either rain or snow) agree with the different water pathways to streams described at other mountainous regions (Brown et al. 2003; Ahearn et al. 2004). On the other hand, air temperature decreased stream discharge of our study sites, suggesting that direct evaporation and plant transpiration across the basin might have a sizable effect on the amount of water reaching the streams. Interestingly, Sosnovsky et al. (2020) reported a positive correlation between air temperature and discharge at shorter timescales during the melting season, suggesting that the influence of air temperature on discharge is both scale and context dependent. Altogether, the same meteorological variable (air temperature) might modulate stream dynamics at several different points along the precipitation/runoff/discharge cycle, sometimes even in opposite direction (e.g., warm temperatures increasing snowmelt but decreasing runoff into stream through evapotranspiration).

Nutrient concentration and annual export rate from CP reflect the usual oligotrophy of north Patagonian streams (Pedrozo et al. 1993; García RD et al. 2015). Values of N and P concentrations were much smaller than those found in either a sub-catchment that had suffered a wildfire (TN = 675.1 µg l^-1^ and TP = 12.0 µg l^−1^) (Temporetti 2006) or in areas of low anthropogenic impact in Patagonia (average TN export = 100 kg km^−2^ yr^−1^) (Little et al. 2008). Indeed, values of P export were between the lowest measured in South America (TP export range from 4 to 450 kg TP km^−2^ yr^−1^) (Álvarez-Cobelas and Angeler 2007). These results indicate that nutrient export in CP stream and similar ecosystems are among the lowest measured for any basin in the Southern hemisphere (Quinn and Stroud 2002; Temporetti 2006; Álvarez-Cobelas and Angeler 2007; Little et al. 2008) and worldwide (Álvarez-Cobelas et al. 2008; Álvarez-Cobelas et al. 2009). Such low export values could be explained by low atmospheric N deposition (Holland 1996) and relatively light land use in this region. It must be noted, however, that nutrient load was measured on a monthly basis, in the absence of precipitation, when overall nutrient export is lower (Abell et al. 2013; Tate and Singer 2013). This implies that our estimates should be considered as baseline values. In any case, it is interesting to note that nutrient export values were higher for G than for CP, even though the opposite result could have been expected, given that headwater lakes act as nutrient and sediment traps (Little et al. 2008; Parker et al. 2009).

Topography and land cover are environmental factors that control spatial patterns in river ecosystems (Snelder and Biggs 2002). For example, the presence of a headwater lake in the drainage basin regulates the flow regime of a stream (Baker et al. 2004). This was made evident by the contrasting values and dynamics of discharge autocorrelation between CP and G streams. The autocorrelation function fell sharply for CP but sloped more gently for G. On the other hand, nutrient exports were higher in G despite the expected trapping effect of its headwater lake. This could be explained by land use of its drainage basin, namely the presence of human settlements and a trout farm. While we did not formally weigh the relative impacts of fish farming and urbanization, we suspect that a growing residential settlement of the Gutiérrez stream drainage basin, often times without adequate wastewater treating infrastructure, is expected to have a larger impact than the detritus from a fish farm producing ∼10 tons of fish per year. Moreover, the additional nutrient sources of fish food and feces from salmonid production would be expected to decrease the TN:TP ratio of the water column in Patagonia reservoirs (Diaz et al. 2001), yet we saw an opposite effect at G stream. Our results suggest that while topography is the main driver of G stream hydrology (Sosnovsky et al. 2020), nutrient dynamics are mainly driven by land use.

The molecular N:P ratio is said to distinguish nitrogen-limited systems from those constrained by phosphorus (Rhee and Gotham 1980). Kahlert (1998, 2001) found that the optimal N:P ratio of freshwater benthic algae was 18:1 (molar), which deviates from the phytoplankton optimal stoichiometric N:P ratio of 16:1 (Redfield ratio) (Goldman et al. 1979). CP and G streams had an arithmetic mean N:P molar ratio close to 15, which would indicate nitrogen deficit in these ecosystems, similar to what has been observed in other ecosystems of Northern Patagonia, either forests (Satti et al. 2003) or lakes (Diaz et al. 2007). However, considering that median values of N:P were 11 for CP and 16 for G, it becomes evident that nitrogen limitation is being lifted at G, most likely due to wastewater from human settlements. Wastewater releases phosphorus and nitrogen to the soil. While Andisols have high capacity in phosphorus retention (Satti et al. 2003), nitrate leaching would reach the stream through the groundwater, priming an eutrophication process of the G stream and downstream water bodies. This finding at the drainage basin level is in line with observations at the global scale suggesting human actions generate a clear unbalance favoring N over P which may induce significant alterations at organism, community and ecosystem levels (Peñuelas et al. 2013).

Leaves of riparian trees are generally the largest component of allochthonous inputs to stream located in forested drainage basins and are often the major source of energy to heterotrophic organisms (Vannote et al. 1980; Webster et al. 1999). In the case of Andean streams, *N. pumilio* leaves cover large areas of the waterway during autumn (Albariño et al. 2009). A substantial amount of these may be transported downstream by spates occurring from late autumn to winter and eventually reaches the lake (Modenutti et al. 2010). We observed no significant differences in the amount of CPOM exported by the two streams studied. However, we highlight the fact that in stream CP the CPOM was mainly made up of native *N. pumilio* leaves, while in stream G introduced *S. fragilis* leaves dominated. This corresponds to the tree species present in the different catchments, and could explain the temporal difference between streams in the surge of CPOM; the peak in CPOM export occurred during June and July in stream CP, and during April in stream G. This supports the claim that afforestation or invasion of exotic plants with different phenology, chemical and physical characteristics to those of the native riparian species changes the timing, quantity and quality of leaf litter standing stock in streams (Naiman et al. 2005).

By studying two Andean streams located at adjacent, yet contrasting drainage basins, we were able to investigate the influence of environmental factors acting at different spatial scales over these freshwater ecosystems. At a regional scale, climate determined seasonality of water flow for both streams. At a landscape scale, we found that the presence of a large, deep lake buffered the hydrological dynamic of its effluent stream. At a lower spatial scale, we found that more intense human impact at G’s drainage basin correlated with higher nutrient export rates by the stream. We could propose, therefore, that the hydrological and nutrients dynamics are decoupled in this ecosystem. Considering the strong demographic pressure at G’s drainage basin, there is a high risk of change in N:P stoichiometry and ecosystem eutrophication. Gutiérrez basin situation thus shows a stark contrast to Casa de Piedra basin, which instead has the hydrological, physical and chemical dynamics of a pristine drainage basin characteristics of Andean streams originating at high altitude wetlands and lakes. Besides geomorphology, the main differences between both basins are land use policies: CP’s basin is mostly within protected areas, while most of G’s lower basin is within the limits of San Carlos de Bariloche city and subject to several, potentially high impact land uses, like residential neighbourhoods, winter sports and fish farming, among others. Local and global environmental problems are currently burning issues in stream ecosystem processes and will be even more so in the future. Considering that lakes and streams are sentinels and integrators of environmental change in the surrounding terrestrial landscape (Williamson et al. 2008), it is really important to implement monitoring schemes for key variables of these aquatic ecosystems (Lovett et al. 2007) and encourage restoration and sustainable management of their drainage basins.

## ACKNOWLEDGMENTS

We thank Rio Negro’s province Water Department and Argentina’s National Weather Service for providing hydrological and meteorological data; and Ahearn D.S. for his interesting comments during the data analysis and writing periods. This study was supported by the Agencia Nacional de Promoción Científica y Tecnológica under Grant number 2959.

Appendix S1. Cross-correlation between climatic variables and discharge (* P ≤ 0.05; * * P ≤ 0.01; * ** P ≤ 0.001; ns: not significant).

| Lag<br>(days) | Discharge-Precipitation |  |  |  | Lag<br>(days) | Discharge-Temperature |  |  |  |
| --- | --- | --- | --- | --- | --- | --- | --- | --- | --- |
| | $r_{CP}$ | $P_{CP}$ | $r_G$ | $P_G$ | | $r_{CP}$ | $P_{CP}$ | $r_G$ | $P_G$ |
| 0 | 0.31 | *** | -0.04 | NS | 0 | -0.38 | *** | -0.13 | ** |
| 1 | 0.30 | *** | -0.03 | NS | 1 | -0.37 | *** | -0.13 | ** |
| 2 | 0.27 | *** | -0.03 | NS | 2 | -0.38 | *** | -0.14 | *** |
| 3 | 0.25 | *** | -0.02 | NS | 3 | -0.39 | *** | -0.15 | *** |
| 4 | 0.26 | *** | -0.01 | NS | 4 | -0.40 | *** | -0.16 | *** |
| 5 | 0.26 | *** | 0.01 | NS | 5 | -0.42 | *** | -0.17 | *** |
| 6 | 0.25 | *** | 0.01 | NS | 6 | -0.44 | *** | -0.19 | *** |
| 7 | 0.23 | *** | 0.01 | NS | 7 | -0.44 | *** | -0.20 | *** |
| 8 | 0.24 | *** | 0.02 | NS | 8 | -0.44 | *** | -0.20 | *** |
| 9 | 0.23 | *** | 0.02 | NS | 9 | -0.44 | *** | -0.21 | *** |
| 10 | 0.19 | *** | 0.01 | NS | 10 | -0.44 | *** | -0.22 | *** |
| 11 | 0.15 | *** | 0.00 | NS | 11 | -0.44 | *** | -0.22 | *** |
| 12 | 0.13 | ** | -0.01 | NS | 12 | -0.46 | *** | -0.24 | *** |
| 13 | 0.14 | ** | -0.00 | NS | 13 | -0.49 | *** | -0.25 | *** |
| 14 | 0.14 | ** | -0.00 | NS | 14 | -0.51 | *** | -0.27 | *** |
| 15 | 0.14 | ** | 0.00 | NS | 15 | -0.53 | *** | -0.29 | *** |
| 16 | 0.12 | ** | -0.00 | NS | 16 | -0.55 | *** | -0.31 | *** |
| 17 | 0.13 | ** | 0.00 | NS | 17 | -0.55 | *** | -0.32 | *** |
| 18 | 0.11 | ** | 0.00 | NS | 18 | -0.55 | *** | -0.34 | *** |
| 19 | 0.10 | ** | -0.00 | NS | 19 | -0.59 | *** | -0.35 | *** |
| 20 | 0.11 | ** | 0.01 | NS | 20 | -0.57 | *** | -0.36 | *** |
| 21 | 0.14 | ** | 0.02 | NS | 21 | -0.57 | *** | -0.36 | *** |
| 22 | 0.16 | ** | 0.03 | NS | 22 | -0.57 | *** | -0.37 | *** |
| 23 | 0.17 | ** | 0.03 | NS | 23 | -0.57 | *** | -0.38 | *** |
| 24 | 0.17 | ** | 0.03 | NS | 24 | -0.57 | *** | -0.39 | *** |
| 25 | 0.19 | *** | 0.05 | NS | 25 | -0.58 | *** | -0.40 | *** |
| 26 | 0.20 | *** | 0.06 | NS | 26 | -0.59 | *** | -0.41 | *** |
| 27 | 0.19 | *** | 0.06 | NS | 27 | -0.60 | *** | -0.43 | *** |
| 28 | 0.20 | *** | 0.07 | NS | 28 | -0.61 | *** | -0.44 | *** |
| 29 | 0.20 | *** | 0.08 | NS | 29 | -0.61 | *** | -0.45 | *** |
| 30 | 0.22 | *** | 0.10 | * | 30 | -0.61 | *** | -0.45 | *** |
| 31 | 0.22 | *** | 0.11 | * | 31 | -0.61 | *** | -0.46 | *** |
| 32 | 0.23 | *** | 0.11 | * | 32 | -0.61 | *** | -0.47 | *** |
| 33 | 0.24 | *** | 0.13 | ** | 33 | -0.60 | *** | -0.48 | *** |
| 34 | 0.24 | *** | 0.14 | ** | 34 | -0.60 | *** | -0.48 | *** |
| 35 | 0.24 | *** | 0.14 | ** | 35 | -0.59 | *** | -0.49 | *** |
| 36 | 0.22 | *** | 0.15 | ** | 36 | -0.59 | *** | -0.49 | *** |
| 37 | 0.20 | ** | 0.14 | ** | 37 | -0.58 | ** | -0.50 | *** |
| 38 | 0.18 | ** | 0.14 | ** | 38 | -0.58 | ** | -0.51 | *** |
| 39 | 0.15 | * | 0.13 | ** | 39 | -0.57 | ** | -0.51 | *** |
| 40 | 0.14 | * | 0.13 | ** | 40 | -0.58 | ** | -0.52 | *** |
| 41 | 0.11 | NS | 0.12 | * | 41 | -0.58 | ** | -0.52 | *** |
| 42 | 0.10 | NS | 0.12 | * | 42 | -0.58 | ** | -0.53 | *** |
| 43 | 0.09 | NS | 0.12 | * | 43 | -0.58 | ** | -0.54 | *** |
| 44 | 0.09 | NS | 0.12 | * | 44 | -0.59 | ** | -0.55 | *** |
| 45 | 0.10 | NS | 0.12 | * | 45 | -0.60 | ** | -0.56 | *** |
| 46 | 0.11 | NS | 0.12 | * | 46 | -0.61 | ** | -0.58 | *** |
| 47 | 0.10 | NS | 0.11 | * | 47 | -0.62 | ** | -0.59 | *** |
| 48 | 0.10 | NS | 0.10 | NS | 48 | -0.64 | ** | -0.60 | *** |
| 49 | 0.11 | NS | 0.11 | * | 49 | -0.66 | ** | -0.62 | *** |
| 50 | 0.13 | NS | 0.12 | * | 50 | -0.67 | ** | -0.63 | *** |
| 51 | 0.12 | NS | 0.11 | NS | 51 | -0.67 | ** | -0.64 | *** |
| 52 | 0.13 | NS | 0.10 | NS | 52 | -0.68 | ** | -0.65 | *** |
| 53 | 0.13 | NS | 0.00 | NS | 53 | -0.68 | ** | -0.66 | *** |
| 54 | 0.16 | NS | 0.00 | NS | 54 | -0.67 | ** | -0.66 | *** |
| 55 | 0.17 | * | 0.00 | NS | 55 | -0.66 | ** | -0.67 | *** |
| 56 | 0.18 | * | 0.00 | NS | 56 | -0.66 | ** | -0.67 | *** |
| 57 | 0.20 | * | 0.00 | * | 57 | -0.66 | ** | -0.68 | *** |
| 58 | 0.23 | ** | 0.00 | * | 58 | -0.67 | ** | -0.69 | *** |
| 59 | 0.25 | ** | 0.00 | * | 59 | -0.68 | ** | -0.69 | *** |
| 60 | 0.24 | ** | 0.00 | * | 60 | -0.68 | ** | -0.70 | *** |
| 61 | 0.25 | ** | 0.00 | ** | 61 | -0.67 | ** | -0.69 | ** |
| 62 | 0.25 | ** | 0.00 | ** | 62 | -0.66 | ** | -0.69 | ** |
| 63 | 0.24 | ** | 0.00 | ** | 63 | -0.66 | ** | -0.69 | ** |
| 64 | 0.23 | * | 0.00 | ** | 64 | -0.67 | ** | -0.69 | ** |
| 65 | 0.23 | * | 0.00 | ** | 65 | -0.67 | ** | -0.69 | ** |
| 66 | 0.23 | * | 0.00 | ** | 66 | -0.65 | ** | -0.69 | ** |
| 67 | 0.24 | ** | 0.00 | ** | 67 | -0.64 | ** | -0.69 | ** |
| 68 | 0.27 | ** | 0.00 | ** | 68 | -0.63 | ** | -0.68 | ** |
| 69 | 0.30 | ** | 0.00 | *** | 69 | -0.63 | ** | -0.68 | ** |
| 70 | 0.31 | ** | 0.00 | *** | 70 | -0.63 | ** | -0.68 | ** |
| 71 | 0.33 | *** | 0.00 | *** | 71 | -0.64 | ** | -0.69 | ** |
| 72 | 0.33 | *** | 0.00 | *** | 72 | -0.64 | * | -0.69 | ** |
| 73 | 0.33 | *** | 0.00 | *** | 73 | -0.64 | * | -0.69 | ** |
| 74 | 0.32 | ** | 0.00 | *** | 74 | -0.63 | * | -0.69 | * |
| 75 | 0.32 | ** | 0.00 | *** | 75 | -0.62 | * | -0.69 | * |
| 76 | 0.31 | ** | 0.00 | *** | 76 | -0.61 | * | -0.69 | * |
| 77 | 0.30 | ** | 0.00 | *** | 77 | -0.62 | * | -0.69 | * |
| 78 | 0.27 | ** | 0.00 | *** | 78 | -0.62 | * | -0.70 | * |
| 79 | 0.25 | * | 0.00 | *** | 79 | -0.62 | * | -0.70 | * |
| 80 | 0.26 | * | 0.00 | *** | 80 | -0.63 | * | -0.71 | * |
| 81 | 0.25 | * | 0.00 | *** | 81 | -0.64 | * | -0.72 | * |
| 82 | 0.26 | * | 0.00 | *** | 82 | -0.65 | * | -0.73 | * |
| 83 | 0.30 | ** | 0.00 | *** | 83 | -0.65 | * | -0.73 | * |
| 84 | 0.30 | ** | 0.00 | *** | 84 | -0.65 | * | -0.73 | * |
| 85 | 0.30 | ** | 0.00 | *** | 85 | -0.64 | * | -0.73 | * |
| 86 | 0.29 | ** | 0.00 | *** | 86 | -0.62 | * | -0.72 | * |
| 87 | 0.29 | ** | 0.00 | *** | 87 | -0.60 | * | -0.71 | * |
| 88 | 0.29 | ** | 0.00 | *** | 88 | -0.59 | * | -0.71 | * |
| 89 | 0.30 | ** | 0.00 | *** | 89 | -0.58 | * | -0.70 | * |
| 90 | 0.30 | ** | 0.00 | *** | 90 | -0.57 | * | -0.69 | * |
| 91 | 0.30 | ** | 0.00 | *** | 91 | -0.55 | * | -0.68 | NS |
| 92 | 0.30 | ** | 0.00 | *** | 92 | -0.54 | * | -0.67 | NS |
| 93 | 0.31 | ** | 0.00 | *** | 93 | -0.53 | NS | -0.67 | NS |
| 94 | 0.31 | ** | 0.00 | *** | 94 | -0.50 | NS | -0.65 | NS |
| 95 | 0.30 | ** | 0.00 | *** | 95 | -0.47 | NS | -0.64 | NS |
| 96 | 0.31 | ** | 0.00 | *** | 96 | -0.45 | NS | -0.62 | NS |
| 97 | 0.33 | ** | 0.00 | *** | 97 | -0.43 | NS | -0.61 | NS |
| 98 | 0.34 | ** | 0.00 | *** | 98 | -0.41 | NS | -0.60 | NS |
| 99 | 0.35 | ** | 0.00 | *** | 99 | -0.40 | NS | -0.59 | NS |
| 100 | 0.37 | ** | 0.00 | *** | 100 | -0.40 | NS | -0.58 | NS |
| 101 | 0.37 | ** | 0.00 | *** | 101 | -0.41 | NS | -0.58 | NS |
| 102 | 0.38 | ** | 0.00 | *** | 102 | -0.41 | NS | -0.57 | NS |
| 103 | 0.37 | ** | 0.00 | *** | 103 | -0.41 | NS | -0.57 | NS |
| 104 | 0.34 | ** | 0.00 | *** | 104 | -0.40 | NS | -0.56 | NS |
| 105 | 0.34 | ** | 0.00 | *** | 105 | -0.40 | NS | -0.55 | NS |
| 106 | 0.33 | ** | 0.00 | *** | 106 | -0.39 | NS | -0.55 | NS |
| 107 | 0.31 | ** | 0.00 | *** | 107 | -0.38 | NS | -0.54 | NS |
| 108 | 0.30 | * | 0.00 | *** | 108 | -0.38 | NS | -0.54 | NS |
| 109 | 0.29 | * | 0.00 | *** | 109 | -0.37 | NS | -0.53 | NS |
| 110 | 0.28 | * | 0.41 | *** | 110 | -0.35 | NS | -0.52 | NS |
| 111 | 0.27 | * | 0.41 | *** | 111 | -0.34 | NS | -0.51 | NS |
| 112 | 0.26 | * | 0.40 | ** | 112 | -0.33 | NS | -0.51 | NS |
| 113 | 0.24 | * | 0.39 | ** | 113 | -0.33 | NS | -0.50 | NS |
| 114 | 0.22 | NS | 0.38 | ** | 114 | -0.33 | NS | -0.50 | NS |
| 115 | 0.21 | NS | 0.37 | ** | 115 | -0.33 | NS | -0.50 | NS |
| 116 | 0.22 | NS | 0.36 | ** | 116 | -0.34 | NS | -0.51 | NS |
| 117 | 0.21 | NS | 0.35 | ** | 117 | -0.34 | NS | -0.51 | NS |
| 118 | 0.22 | NS | 0.35 | ** | 118 | -0.34 | NS | -0.51 | NS |
| 119 | 0.23 | * | 0.35 | ** | 119 | -0.32 | NS | -0.50 | NS |
| 120 | 0.24 | * | 0.35 | ** | 120 | -0.29 | NS | -0.49 | NS |
| 121 | 0.22 | NS | 0.34 | ** | 121 | -0.28 | NS | -0.48 | NS |
| 122 | 0.24 | * | 0.34 | * | 122 | -0.27 | NS | -0.48 | NS |
| 123 | 0.24 | * | 0.34 | * | 123 | -0.25 | NS | -0.47 | NS |
| 124 | 0.26 | * | 0.34 | * | 124 | -0.24 | NS | -0.46 | NS |
| 125 | 0.25 | * | 0.34 | * | 125 | -0.22 | NS | -0.45 | NS |
| 126 | 0.23 | NS | 0.33 | * | 126 | -0.21 | NS | -0.43 | NS |
| 127 | 0.25 | * | 0.35 | * | 127 | -0.20 | NS | -0.43 | NS |
| 128 | 0.24 | * | 0.35 | * | 128 | -0.21 | NS | -0.42 | NS |
| 129 | 0.25 | * | 0.36 | ** | 129 | -0.21 | NS | -0.41 | NS |
| 130 | 0.25 | * | 0.36 | ** | 130 | -0.21 | NS | -0.40 | NS |
| 131 | 0.25 | * | 0.36 | ** | 131 | -0.20 | NS | -0.40 | NS |
| 132 | 0.24 | * | 0.36 | ** | 132 | -0.19 | NS | -0.39 | NS |
| 133 | 0.26 | * | 0.38 | ** | 133 | -0.17 | NS | -0.38 | NS |
| 134 | 0.25 | * | 0.38 | ** | 134 | -0.15 | NS | -0.36 | NS |
| 135 | 0.27 | * | 0.39 | ** | 135 | -0.12 | NS | -0.35 | NS |
| 136 | 0.27 | * | 0.39 | ** | 136 | -0.10 | NS | -0.33 | NS |
| 137 | 0.25 | * | 0.38 | ** | 137 | -0.08 | NS | -0.32 | NS |
| 138 | 0.26 | * | 0.37 | * | 138 | -0.06 | NS | -0.30 | NS |
| 139 | 0.26 | * | 0.37 | * | 139 | -0.06 | NS | -0.29 | NS |
| 140 | 0.26 | * | 0.37 | * | 140 | -0.06 | NS | -0.28 | NS |
| 141 | 0.25 | * | 0.36 | * | 141 | -0.06 | NS | -0.27 | NS |
| 142 | 0.24 | NS | 0.35 | * | 142 | -0.04 | NS | -0.25 | NS |
| 143 | 0.22 | NS | 0.35 | * | 143 | -0.01 | NS | -0.23 | NS |
| 144 | 0.20 | NS | 0.33 | * | 144 | 0.02 | NS | -0.21 | NS |
| 145 | 0.15 | NS | 0.31 | * | 145 | 0.04 | NS | -0.20 | NS |

## REFERENCES

A.P.H.A. 2005. Standard methods for examination of water and wastewater. 21 ed. Washington DC: American Public Health Association.

Abell JM, Hamilton DP, Rutherfor JC. 2013. Quantifying temporal and spatial variations in sediment, nitrogen and phosphorus transport in stream inflows to a large eutrophic lake. Envirnomental Science Processes and Impacts.

Ahearn DS, Sheibley RW, Dahlgren RA, Keller KE. 2004. Temporal dynamics of stream water chemistry in the last free-flowing river draining the western Sierra Nevada, California. Journal of Hydrology. 295:47–63.

Albariño RJ, Díaz Villanueva V, Buria L. 2009. Leaf litter dynamics in a forested small Andean catchment, northern Patagonia, Argentina. In: Oyarzún C, Verhoest NEC, Boeckx P et al., editors. Ecological advances on Chilean temperate rainforest. Academia Press; p. 183–211.

Álvarez-Cobelas M, Angeler DG. 2007. Exportación de nutrientes en las cuencas hidrográficas de Latinoamérica: una recopilación. Revista Latinoamericana de Recursos Naturales. 3(1):31–43.

Álvarez-Cobelas M, Angeler DG, Sánchez-Carrillo S. 2008. Export of nitrogen from catchments: A worldwide analysis. environmental Pollution. 156:261–269.

Álvarez-Cobelas M, Sánchez-Carrillo S, Angeler DG. 2009. Phosphorus export from catchments: a global view. Journal of American Benthological Society. 28(4):805–820.

Allan JD. 2004. Landscapes and Riverscapes: The influence of land use on stream ecosystems. Annual Review of Ecology, Evolution and Systematics. 35:257–284.

Bailey RG. 1995. Ecosystem Geography. New York, New York: Springer-Verlag.

Baker DB, Richards RP, Loftus TT, Kramer JW. 2004. A new flashiness index: Characteristics and applications to midwestern rivers and streams. Journal of the American Water Resources Association. 40(2):503–522.

Brown LE, Hannah DM, Milner AM. 2003. Alpine Stream Habitat Classification: An Alternative Approach Incorporating the Role of Dynamic Water Source Contributions. Artic, Antartic, and Alpine Research. 35(3):313–322.

Carpenter SR, Caraco NF, Correll DL, Howarth RW, Sharpley AN, Smith VH. 1998. Nonpoint Pollution of Surface Waters with Phosphorus and Nitrogen. Ecological Applications. 8(3):559–568.

Carrillo U, Díaz Villanueva V, Modenutti BE. 2018. Sustained effects of volcanic ash on biofilm stoichiometry, enzyme activity and community composition in North-Patagonia streams. Science of the Total Environment. 621:235–244.

Diaz MM, Pedrozo FL, Reynolds CS, Temporetti PF. 2007. Chemical composition and the nitrogen-regulated trophic state of Patagonian lakes. Limnologica. 37:17–27.

Diaz MM, Temporetti PF, Pedrozo FL. 2001. Response of phytoplankton to enrichment from cage fish farm waste in Alicura Reservoir (Patagonia, Argentina). Lake & Reservoir Management. 6:151–158.

Díaz Villanueva V, Bastidas Navarro M, Albariño R. 2016. Seasonal patterns of organic matter stoichiometry along a mountain catchment. Hydrobiologia.

Diehl L, Mazzarino MJ, Funes F, Fontenla S, Gobbi M, Ferrari J. 2003. Nutrient conservation strategies in native Andean-Patagonian forests. Journal of Vegetation Science. 14:63–70.

Dodds WK, Smith VH. 2016. Nitrogen, phosphorus, and eutrophication in streams. Inland Waters. 6:155–164.

Emerson DG, Vecchia AV, Dahl AL. 2005. Evaluation of Drainage-Area Ratio Method Used to Estimate Streamflow for the Red River of the North Basin, North Dakota and Minnesota. Reston, Virginia: U.S. Geological Survey.

García DR, Diéguez MC, Gerea M, García PE. 2018. Characterisation and reactivty continuum of dissolved organic matter in forested headwater catchments of Andean Patagonia. Freshwater Biology. 63(9):1049–1062.

García P, Diéguez MC, Queimaliños CP. 2015. Landscape integration of North Patagonian mountain lakes: a first approach using characterization of dissolved organic matter. Lakes & Reservoirs: Research and Management. 20:1–14.

García RD, Reissig M, Queimaliños CP, García PE, Diéguez MC. 2015. Climate-driven terrestrial inputs in ultraoligotrophic mountain streams of Andean Patagonia revealed through chromophoric and fluorescent dissolved organic matter. Science of the Total Environment. 521–522:280–292.

Giacosa R, Heredia N, González R, Faroux A, Césari O, Franchi M. 2001. Hoja Geológica 4172-IV, San Carlos de Bariloche. Provincias de Río Negro y Neuquén.

Goldman JC, Mc Carthy JJ, Peavey DG. 1979. Growth rate influence on the chemical composition of phytoplankton in oceanic waters. Nature. 279:210–215.

Gordon ND, McMahon TA, Gippel CJ, Natha RJ. 2004. Stream Hydrology An Introduction for Ecologist. Chichester, England: John Wiley & Sons LTD.

Grasshoff M, Ehrhardt K, Kremling K. 1983. Methods of seawater analysis. New York: Wiley.

Hintze JL. 1998. Number Cruncher Statistical System (NCSS). Version 2000. User’s Guide. Utah: Kaysville.

Holland EA, Dentener FJ, Braswell BH, Sulzman JM. 1999. Contemporary and pre-industrial global reactive nitrogen budgets. Biogeochemistry. 46:7–43.

Howarth RW, Billen G, Swaney D, Townsend A, Jaworski NA, Lajtha K, Downing JA, Elmgren R, Caraco N, Jordan T et al. 1996. Regional nitrogen budgets and riverine N & P fluxes for the drainages to the North Atlantic Ocean: natural and human influences. Biogeochemistry. 35:75–139.

Hynes HBN. 1975. The stream and its valley. Verhandlungen der Internationale Vereingung für Limnologie. 19:1–15.

Jordan TE, Correll DL, Weller DE. 1997. Effects of agriculture on discharges of nutrients from coastal plain watersheds of Chesapeake Bay. Journal of Environmental Quality. 26:836–848.

Kahlert M. 1998. C:N:P ratios of freshwater benthic algae. Archiv für Hydrobiologie, Special Issues, Advances in Limnology. 51:105–114.

Kahlert M. 2001. Biomass and Nutrient Status of Benthic Algae in Lakes. Acta Universitatis Upsaliensis: Uppsala.

Kalff J. 2002. Chapter 5, Hidrology and Climate. In: Ryu T, editor. Limnology Inland Water Ecosystems. Upper Saddle River: Prentice Hall; p. 53–71.

Lallement M, Macchi PJ, Vigliano PH, Juarez S, Rechencq M, Baker M, Bouwes N, Crowl TA. 2016. Rising from the ashes: Change in salmonid fish assemblages after 30 months of the Puyehue-Cordon Caulle volcanic eruption. Science of the Total Environment. 541:1041–1051.

Larocque M, Mangin A, Razack M, Banton O. 1998. Contribution of correlation and spectral analyses to the regional study of a large karst aquifer (Charente, France). Journal of Hydrology. 205:217–231.

Little C, Soto D, Lara A, Cuevas JG. 2008. Nitrogen exports at multiple-scales in a southern Chilean watershed (Patagonian Lakes district). Biogeochemistry. 87:297–309.

Lovett GM, Burns DA, Driscoll CT, Jenkins JC, Mitchell MJ, Rustad L, Shanley JB, Likens GE, Haeuber R. 2007. Who need environmental monitoring? Frontiers in Ecology and the Envirnonment. 5(5):253–260.

Mangin A. 1984. Pour une meilleure connaissance des systèmes hydrologgiques à partir des analyses corrélatoire et spectrale. Journal of Hydrology. 67:25–43.

Masiokas MH, Villalba R, Luckman BH, Lascano ME, Delgado S, Stepanek P. 2008. 20th-century recession and regional hydroclimatic changes in northwestern Patagonia. Global and Planetary Change. 60:85–100.

Merseburger GC, Martí E, Sabater F. 2005. Net changes in nutrient concentrations below a point source input in two streams draining catchments with contrasting land uses. Science of the Total Environment. 347:217–229.

Meyer JL, Paul MJ, Taulbee WK. 2005. Stream ecosystem function in urbanizing landscapes. Journal of the North American Benthological Society. 24:602–612.

Miserendino ML, Kutschker MA, Brand C, La Manna L, Di Prinzio YC, Papazian G, Bava J. 2016. Ecological Status of a Patagonian Mountain River: Usefulness of Environmental and Biotic Metrics for Rehabilitation Assessment. Environmental Management. 57(6):1166–1187.

Modenutti BE, Albariño RJ, Bastidas Navarro M, Villanueva VD, Souza AF, Trochine C, Laspoumaderes C, Cuassolo F, Mariluán G, Buria L et al. 2010. Structure and dynamic of food webs in Andean North Patagonian freshwater systems: organic matter, light and nutrient relationships. Ecología Austral. 20(2):95–114.

Naiman RJ, Décamps H, McClain ME. 2005. Riparia: ecology, conservation and management of streamside communities. London, UK: Elsevier Academic Press.

Oyarzún CE, Godoy R, Schrijver De A, Staelens J, Lust N. 2004. Water chemistry and nutrient budgets in an undisturbed evergreen rainforest of souther Chile. Biogeochemistry. 71:107–123.

Parker BP, Schindler DE, Beaty KG, Stainton MP, Kasian SEM. 2009. Long-term changes in climate, streamflow, and nutrient budgets for first-order catchments at the Experimental Lakes Area (Ontario, Canada). Canadian Journal of Fisheries and Aquatic Sciences. 66:1848–1863.

Paruelo JM, Beltran A, Jobbágy E, Sala O, Golluscio R. 1998. The climate of Patagonia: general patterns and controls on biotic processes. Ecología Austral. 8(2):85–101.

Pedrozo FL, Chillrud S, Temporetti PF, Diaz MM. 1993. Chemical composition and nutrient limitation in rivers and lakes of northern Patagonian Andes (39.5º-42º S; 71º W) (Rep. Argentina). SIL Proceedings, 1922–2010. 25:207–214.

Peel MC, Finlayson BL, McMahon TA. 2007. Updated world map of the Köppen-Geiger climate classification. Hydrology and Earth System Sciences. 4(2):439–473.

Peltier MR, Wilcox CJ, Sharp DC. 1998. Technical Note: Application of the Box-Cox data transformation to animal science experiments. Journal of Animal Science. 78:847–849.

Peñuelas J, Poulter B, Sardans J, Ciais P, van der Velde M, Bopp L, Boucher O, Godderis Y, Hinsinger P, Llusia J et al. 2013. Human-induced nitrogen– phosphorus imbalances alter natural and managed ecosystems across the globe. Nature Communications. 4:2934.

Pereyra FX, Albertoni J, Bréard C, Cavaliaro S, Coccia M, Ducós E, Dzendoletas M, Fookes S, Getino E, Helms F et al. 2005. Estudio geocientífico aplicado al ordenamiento territorial, San Carlos de Bariloche.

Pieterse NM, Bleuten W, Jørgensen SE. 2003. Contribution of point sources and diffuse sources to nitrogen and phosphorus loads in lowland rivers tributaries. Journal of Hydrology. 271:213–225.

Poff NL, Allan JD, Bain MB, Karr JR, Prestegaard KL, Richter BD, Sparks RE, Stromberg JC. 1997. The Natural Flow Regime. Bioscience. 47(11):769–784.

Pozo J, Elosegi A, Díez J, Molinero J. 2009. Chapter 10, Dinámica y relevancia de la materia orgánica. In: Elosegi A, Sabater S, editors. Conceptos y técnicas en ecología fluvial. Bilbao, España: Fundación BBVA; p. 141–166.

Quinn JM, Stroud MJ. 2002. Water quality and sediment and nutrient export from New Zealand hill-land. New Zealand Journal of Marine and Freshwater Research. 36(2):409–429.

Rhee GY, Gotham IJ. 1980. Optimum N:P ratios and co-existence of planktonic algae. Journal of Phycology. 16:486–489.

Satti P, Mazzarino MJ, Gobbi M. 2003. Soil N dynamics in relation to leaf litter quality and soil fertility in north-western Patagonian forests. Journal of Ecology. 91:173–181.

Snelder TH, Biggs BJF. 2002. Multiscale river environment classification for water resources management. Journal of the American Water Resources Association. 38:1225–1239.

Sosnovsky A, Rechencq M, Fernández V, Suarez MJ, Cantet RJC. 2020. Hydrological and Physico-chemical dynamics in two Andean streams. Limnetica. 39(1):17–33.

Tate KW, Singer MJ. 2013. Timing, Frecuency of Sampling Affect Accuracy of Water-Quality Monitoring. California Agriculture. 53(6):44–48.

Temporetti PF. 2006. Efecto a largo plazo de los incendios forestales en la calidad del agua de dos arroyos en la sub-región Andino-Patagónica, Argentina. Ecología Austral. 16(2):157–166.

Vannote RL, Minshall GW, Cummins KW, Sedell JR, Cushing CE. 1980. The River Continuum Concept. Canadian Journal of Fisheries and Aquatic Sciences. 37:130–137.

Walsh CJ, Fletcher TD, Ladson AR. 2005. Stream restoration in urban catchments through redesigning stormwater systems: looking to the catchment to save the stream. Journal of the North American Benthological Society. 24(690–705).

Webster JR, Benfield EF, Ehrman TP, Schaeffer MA, Tank JL, Hutchens JJ, D’angelo DJ. 1999. What happens to allochthonous material that falls into streams? A synthesis of new and published information from Coweeta. Freshwater Biology. 41:687–705.

Wiens JA. 2002. Riverine landscapes: taking landscape ecology into the water. Freshwater Biology. 47:501–515.

Williams Subiza EA, Brand C. 2018. Short-term effect of wildfire on Patagonian headwater streams. International Journal of Wildland Fire. 27:457–470.

Williamson CE, Dodds WK, Kratz TK, Palmer MA. 2008. Lakes and streams as sentinels of environmental change in terrestrial and atmospheric processes. Frontiers in Ecology and the Envirnonment. 6(5):247–254.

